# Long-term fungus–plant co-variation from multi-site sedimentary ancient DNA metabarcoding in Siberia

**DOI:** 10.1101/2021.11.05.465756

**Authors:** Barbara von Hippel, Kathleen R. Stoof-Leichsenring, Luise Schulte, Peter Seeber, Laura S. Epp, Boris K. Biskaborn, Bernhard Diekmann, Martin Melles, Luidmila Pestryakova, Ulrike Herzschuh

**Author notes:** <u>corresponding author</u>.

## Abstract

Climate change has a major impact on arctic and boreal terrestrial ecosystems as warming leads to northward treeline shifts, inducing consequences for heterotrophic organisms associated with the plant taxa. To unravel ecological dependencies, we address how long-term climatic changes have shaped the palaeo-ecosystems at selected sites in Siberia.

We investigated sedimentary ancient DNA from five lakes spanning the last 47,000 years, using the ITS1 marker for fungi and the chloroplast P6 loop marker for vegetation metabarcoding. After bioinformatic processing with the OBItools pipeline, we obtained 706 unique fungal operational taxonomic units (OTUs) and 243 amplicon sequence variants (ASVs) for the plants. We show higher OTU numbers in dry forest tundra as well as boreal forests compared to wet southern tundra. The most abundant fungal taxa in our dataset are Pseudeurotiaceae, *Mortierella*, Sordariomyceta, *Exophiala, Oidiodendron, Protoventuria, Candida vartiovaarae, Pseudeurotium, Gryganskiella fimbricystis*, and *Trichosporiella cerebriformis*. The overall fungal composition is explained by the plant composition as revealed by redundancy analysis. The fungal functional groups show antagonistic relationships in their climate susceptibility. The advance of woody taxa in response to past warming led to an increase in the abundance of mycorrhizae, lichens, and parasites, while yeast and saprotroph distribution declined. We also show co-occurrences between Salicaceae, *Larix*, and *Alnus* and their associated pathogens and detect higher mycorrhizal fungus diversity with the presence of Pinaceae. Under future warming, we can expect feedbacks between fungus compositional and plant diversity changes which will affect forest advance and stability in arctic regions.

## 1. Introduction

Fungi represent a principal component of soils and sustain a broad variety of ecosystem functions (Frąc et al., 2018) that are highly dependent on the fungal composition and are subject to changing environmental conditions. For example, experimental warming led to an increase in ectomycorrhizal fungi and free-living filamentous fungi, while a decrease in yeast was observed (Treseder et al., 2016). Besides, experimental warming showed an increase in the evenness of fungal tundra communities, including a significant increase in ectomycorrhizal fungi in relation to rising temperature (Deslippe et al., 2012). Furthermore, under warming, saprotrophs shifted their metabolism from decaying wood to self-maintenance and subsequent spore-production (Romero-Olivares et al., 2015; 2019). It has also been reported that warming leads to an increase in specific plant pathogens (Otrosina and Cobb, 1989). Accordingly, time-series on compositional changes of fungal communities during past climate changes are highly valuable to assess the potential shifts of ecosystem functioning in a rapidly warming world. Time-series are also an asset when testing whether lab experiments reflect real natural developments on long geological timescales.

Fungal functional types reflect the strong interactions between plants and fungi but we are far from understanding the detailed processes involved in particular temporal changes in their covariation (Zobel et al., 2018). These types help group fungi according to their functional roles in the terrestrial ecosystem allowing the assessment of compositional shifts. This, however, remains a challenging task as the ecological functions of many taxa are still not understood. The major ecological functional fungus groups in forest ecosystems are saprotrophs, mycorrhizae, and parasites. Saprotrophic species are major decomposers in terrestrial ecosystems (Baldrian and Valášková, 2008). A plant’s benefit from mycorrhizal fungi is the enabled acquisition of mineral nutrients in solution, such as phosphate, while the fungus in return receives carbohydrates from the plants (Finlay, 2008). Mycorrhizal fungi-plant associations include arbuscular mycorrhizae, ectomycorrhizae, ericoid mycorrhizae, and orchid mycorrhizae (Brundrett and Tedersoo, 2018). Parasitic fungi, such as *Heterobasidion*, infecting conifer tree taxa (Garbelotto and Gonthier, 2013), are important for eliminating weak trees to maintain the functioning of healthy forest ecosystems. Biotrophic plant parasites feed from living tissue while necrotrophic fungi penetrate the plant, destroy the tissue, and subsequently provoke plant death (Naranjo-Ortiz and Gabaldón, 2019).

Boreal forests are the world’s largest biome and cover an area of around 9% of the total land mass between 45° and 70° north (Czimczik, 2005). These ecosystems are an important habitat for fungi and in particular arbuscular mycorrhizal fungi (Öpik et al., 2008). Ectomycorrhizal communities in the sub-arctic tundra are generally species-rich but do not show a high host preference (Ryberg et al., 2009; 2011). Nevertheless, for the establishment of these mycorrhizal associations, fungal preferences towards distinct plant families or even species were detected. For example, *Glomus intraradices* (Smith et al., 1998; Wagg et al., 2008) and also *Suillus* species have been found to predominantly build up mycorrhizal associations with Pinaceae (Palm and Stewart, 1984; Wagg et al., 2008; Liao et al., 2016). Therefore, the ongoing invasion of woody taxa into tundra areas due to global warming should impact fungus composition. Furthermore, deep DNA sequencing studies of soil fungi in arctic Alaska revealed that warming does not affect overall fungus richness but leads to a change in the community composition. The results of these studies revealed that a decrease in ectomycorrhizae, ericoid mycorrhizae, and lichens in the tundra accompanied an increase of saprotrophic, pathogenic, and root endophytic fungal richness (Geml et al. 2015; Mundra et al. 2016). Hitherto almost all information on fungal compositional turnover originates from short-term warming experiments. Therefore, it is necessary to analyse long-term changes in plant-fungi covariation under natural conditions in different settings including different time scales to include potential lags of vegetation responses towards short-term warming.

Compositional changes of vegetation and associated fungal communities are slow. They occur on decadal, centennial, or even millennial time-scales which are not covered by the, anyway, rare observational time-series and thus the exploitation of palaeoecological archives is required. As large parts of Siberia were not covered by glaciers during the Last Glacial Maximum (Svendsen et al., 2004), lakes from this region provide sedimentary archives which continuously cover the rather warm marine isotopic stage (MIS) MIS 3 (50-30 ka), the cold MIS 2 (30-15.5 ka), and the warm Holocene (MIS 1) (the last 11.6 ka) (Kreveld et al., 2000; Swann et al., 2005), thereby encompassing tremendous vegetation changes. Lake sediments represent natural archives of terrestrial environmental change (Courtin et al., 2021). Given that northern Russia is warming faster than the global average (Biskaborn et al. 2019), lake sediments can provide valuable information on the associated terrestrial ecosystem changes. While many sedimentary pollen records focus on vegetation change, there is limited information about fungi (e.g. from non-pollen palynomorphs (Van Geel, 2001)) as they lack a morphological fossil record.

Sedimentary ancient DNA metabarcoding (sedaDNA) is a promising palaeoecological proxy method which makes use of specific genetic marker regions enabling the study of past biodiversity (Sønstebø et al., 2010). So far, many sedaDNA studies have investigated plant metabarcoding, mostly applying the trnL P6 loop marker (Parducci et al., 2017; Alsos et al., 2018; Liu 2020), but there are only few studies focusing on fungal ancient DNA from sedimentary deposits (Lydolph et al., 2005; Bellemain et al., 2013; Talas et al., 2021). Bellemain et al. (2013) traced past fungal communities in Siberian permafrost sediments. In addition to permafrost, lake sediments can also be used to investigate past ecosystem dynamics. Through relocation processes, the environmental DNA is continuously deposited in the lake which favours these sediments over soil profiles. Recently, Talas et al. (2021) investigated lake sediments showing clear variations in the community compositions: while saprotrophs remained stable over time, host-specific fungi such as plankton parasites and mycorrhizae shifted in relation to human impact and changing climate. The internal transcribed spacer (ITS) region is the most commonly used DNA barcoding region for fungi (Seifertt, 2009, but the primers that were used in early studies caused quite some amplification biases (Bellemain et al., 2010). Primers to specifically target ancient and degraded DNA were designed and used on permafrost deposits (Epp et al., 2012; Bellemain et al., 2013). These have now been refined according to the current status of reference databases by Seeber et al. (2021), providing a primer pair that is highly suitable to amplify sedaDNA as it targets short amplicons of a mean length of 183 bp and is highly specific towards fungi. This enables now studies using sedaDNA to trace fungus-plant interactions over time.

The aim of this study is to analyse lake sediments from five sites in Siberia spanning MIS 3, 2, and 1 for their fungal composition using sedaDNA metabarcoding with the ITS1 marker. The obtained data are compared to vegetation data gained from metabarcoding with the trnL P6 loop marker on the same samples. This study addresses the following questions: (1) How does fungal alpha diversity change during tundra-forest transitions? (2) How do fungal taxonomic composition and function change relative to vegetation transition? Based on the answers, we draw conclusions on fungus-plant covariation under climatic changes over long timescales. These results will help to improve predictions of the response of fungus-plant interactions to future warming.

## 2. Geographic setting and study sites

All study sites are located within Siberia, central eastern Russia (Figure 1), and are characterised by permafrost soils (Brown et al. 1997; Tchebakova et al., 2009). The climate in the entire area is rather continental with hot summers and long, severe winters (*Atlas Arktiki*, 1985). The most prevalent vegetation is boreal forest with spruce, pine, fir, and larch in the western and southern parts and pure larch forest in the east. The arctic regions along the coast and on the Taymyr Peninsula are covered by tundra.

**Fig. 1:**
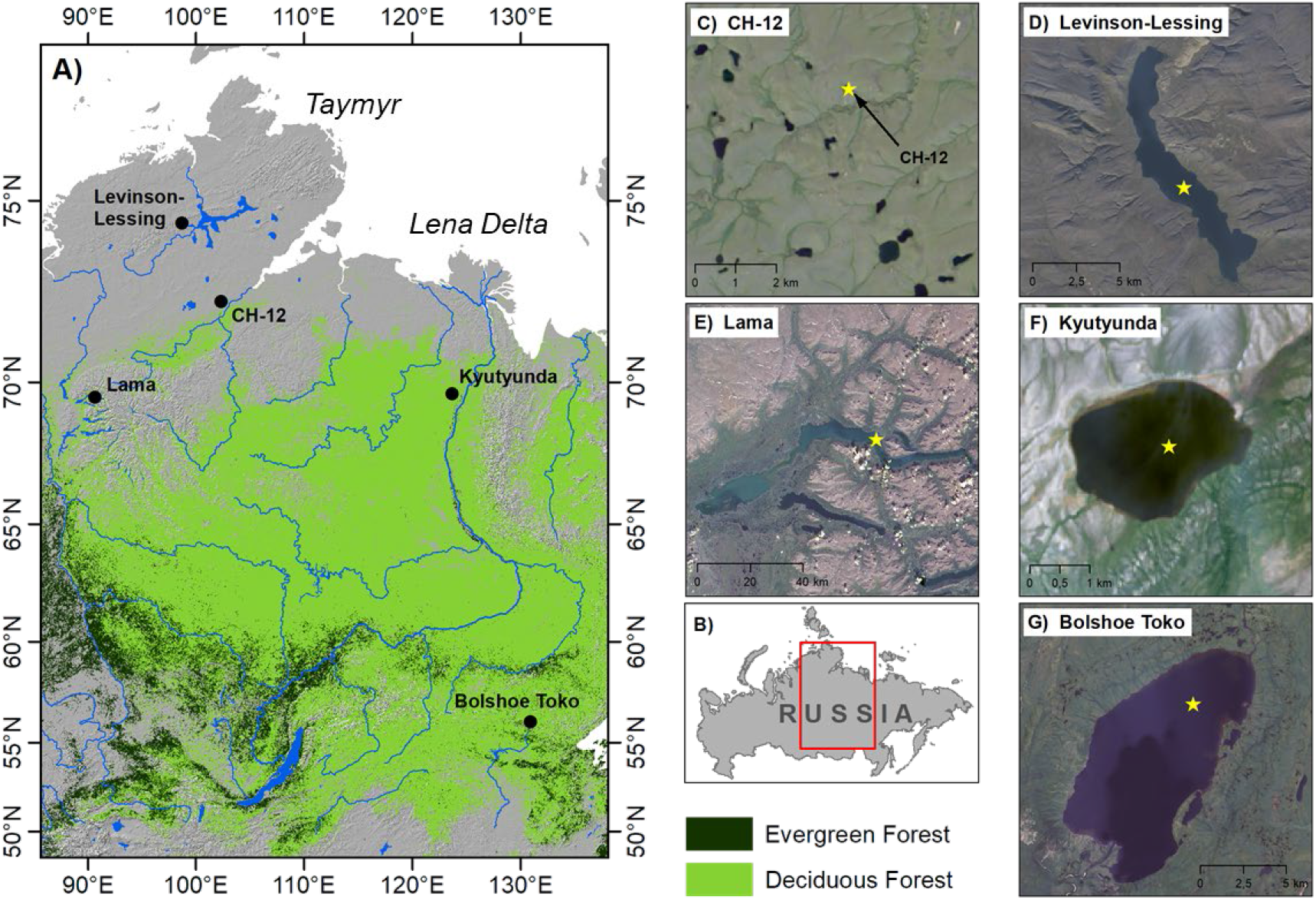
Map of central Russia showing the location of the study sites. (C) to (G) satellite images of the lakes with their surroundings (core names are indicated in brackets). The locations of the cores are marked with an asterisk.

Lake Levinson Lessing (74.27°N, 98.39°E; 48 m a.s.l.) is located in the tundra region of the Taymyr Peninsula. The lake is surrounded by sparse lichen-herb tundra, moss-forb tundra, and dry sedge-forb tundra with dominant *Dryas octopetala, Salix polaris*, and *Cassiope tetragona* (Anisimov and Pospelov, 1999). Mean July temperature is 12.5°C and mean January temperature -31.5 °C (Hatanga weather station; 71.98 °N, 102.47 °E; distance to the lake: 289 km (WorldRussian Institute of Hydrometeorological Information: Data Center, 2021)). The lake is approximately 15 km long and has a width of 2.5 km and a maximum depth of 120 m (Lebas et al., 2019). In an expedition in April 2017, a 46 m sediment core (Co1401; Figure 1D) was retrieved from the central part of the lake at a depth of 112m. According to the chronology, the core covers the last 62 cal ka BP (Scheidt et al., 2021).

Lake CH12 (72.4°N, 102.29°E; 60 m a.s.l.) is located at the tree line on the southern Taymyr Peninsula north of the Putorana Plateau. The current vegetation in the study area is shrub tundra dominated by *Sphagnum, Hylocomium, Aulacomnium, Dicranum*, and *Polytrichum* as well as *Empetrum nigrum, Betula nana*, and *Vaccinium oliginosum*. Also stands of *Larix gmelinii* are present (Klemm et al., 2016; Niemeyer et al., 2017). Mean July temperature is 12.5 °C and the mean January temperature is -31.5 °C (Hatanga weather station; 71.98 °N, 102.47 °E; distance to the lake: 60 km (Russian Institute of Hydrometeorological Information: World Data Center, 2021)). The lake is elliptically shaped with a mean radius of 100 m. The maximum depth of the lake is 14.3 m. A 1.21 m long sediment core was retrieved in an expedition in 2011 (Figure 1C). The core dates back to 7.1 cal ka BP (Stoof-Leichsenring et al., 2015; Klemm et al., 2016).

Lake Kyutyunda (69.38 °N, 123.38°E; 66 m a.s.l.) is located in northern Siberia on the central Siberian Plateau in the tundra-taiga transition zone, which is formed of a mosaic of *Larix* forest and shrub tundra with Poaceae, *Dryas*, and *Saxifraga* species. The data from the Kjusjur weather station (70.68 °N, 127.4 °E; distance to the lake: 210 km (Russian Institute of Hydrometeorological Information: World Data Center, 2021)) give mean July temperatures of 13 °C and mean January temperatures of -35.3 °C. The lake is roughly circular at 2.2 km long by 3 km. The deepest part of this lake is 3.5 m. In an expedition in 2010, a 7 m sediment core (PG2023) dating back to 38.8 cal ka BP was collected (Biskaborn et al., 2016). For the lake Kyutyunda sediment core (Figure 1F) we used an updated version of the age-depth correlation from Biskaborn et al. (2016) with refined correlation between the core segments (Supplementary 1 and 2), based on the IntCal20 calibration curve (Reimer et al., 2020) compiled in the R package bacon (Blaauw and Christen, 2011).

Lake Lama (69.32°N, 90.12°E; 53 m a.s.l.) is located on the Putorana Plateau. Low elevations are covered by dense taiga with *Picea, Larix*, and *Betula*, shrubs such as *Alnus fructicosa, Salix*, and *Juniperus communis*, and dwarf shrubs (Andreev et al., 2004). Modern temperatures vary between a mean of 13.8 °C in July and -28.8 °C in January (Volochanka weather station; 70.97 °N, 94.5 °E ; distance to the lake: 247 km (Russian Institute of Hydrometeorological Information: World Data Center, 2021)). This lake covers an area of 318 km^2^, is 80 km long and up to 7 km wide. It has a maximum depth of 254 m. A sediment core with a total length of 18.85 m (PG1341; Figure 1E) was retrieved from a water depth of 66 m during an expedition in 1997. The core dates back to 23 cal ka BP. The age-depth model of the sediment core is described in this study (Supplementary 3 and 4). Nineteen sediment samples of Lake Lama were radiocarbon dated with an accelerator mass spectrometer (AMS) MICADAS (MIni CArbon Dating System) at Alfred Wegener Institute Bremerhaven and an age-depth model has been established using the R package bacon (Blaauw and Christen, 2011).

Lake Bolshoe Toko (56.15° N, 130.30 °E; 903 m a.s.l.) is located on the northern slope of the eastern Stanovoy Mountain Range in southern central Yakutia in an area with deciduous boreal forests formed by *Larix cajanderi* and *L. gmelinii* with occurrences of *Picea obovata, P. jezoensis*, and *Pinus sylvestris* (Konstantinov 2000). The temperature in the area varies between 34 °C in July to -65 °C in January (Toko weather station; 56.1 °N, 131.01 °E; distance to the lake: 44 km (Konstantinov, 2000)). The lake is 15.4 km long and 7.5 km wide, and has a maximum depth of 72.5 m. In an expedition in 2013 (Biskaborn et al., 2019), a 3.8 m long sediment core (PG2133; Figure 1G) was retrieved from the lake at 26 m water depth. The core dates back to 33.8 cal ka BP (Courtin et al., 2021).

## 3. Materials and Methods

### 3.1 Sampling

SedaDNA samples were taken from 1 m long sub-core segments that were cut in half. Subsampling was undertaken in the climate chamber of the Helmholtz Centre Potsdam – German Research Centre for Geosciences (GFZ) at 10 °C. The chamber is located in the cellar of the institute building in which no molecular genetic studies are conducted. Before subsampling, all surfaces in the chamber were cleaned with DNA Exitus Plus™ (VWR, Germany) and demineralised water. All knives, scalpel holders, and other sampling tools were cleaned according to the recommendations of Champlot et al. (2010) to avoid any contamination with modern DNA and amongst the samples themselves. All materials used for the sampling were taken from the palaeogenetic DNA laboratory at the Alfred Wegener Institute (AWI) in Potsdam where they had been treated to remove DNA.

During the sampling, protective clothing as well as face masks were worn. The surface of the core halves were scraped off twice with sterile scalpel blades and the sample was taken with the help of four knives or two aluminium disks to cut-off the sample and then placed in sterile 8 mL Sarstedt tubes. All samples were taken under the same conditions. The core from Lake Levinson Lessing was sampled under the same conditions in the laboratories of the Institute of Geology and Mineralogy at the University of Cologne.

The samples were analysed according to their estimated ages, at intervals of about 5 cal kyr, leading to 15 samples from lake Lama, 9 samples from lake Levinson Lessing, 10 samples from lake Kyutyunda and and 8 samples from lake Bolshoe Toko. For lake CH12, 28 samples were taken, representing every 100-250 years.

### 3.2 DNA extraction and amplification

SedaDNA was extracted using the DNeasy PowerMax Soil DNA Isolation Kit (Qiagen, Germany) according to the manufacturer’s instructions. An additional incubation step at 56 °C in a rotation oven overnight after mixing the samples with the PowerMax beads was included. Proteinase K (2 mg mL^-1^) and DTT (5 M) were added before the incubation. The final elution step was conducted using 2 mL of the solution C6. Each extraction batch was processed on a different day to avoid contamination between the batches. The 70 samples were investigated for both fungi and vegetation. A volume of 0.5 mL of the CH12 extracts was purified and at the same time concentrated to 50 μL with a GeneJET PCR purification Kit (Thermo Fisher Scientific, Germany). For the other lakes, 1 mL of the DNA extract was used for the purification. The DNA concentration of concentrated samples was measured with a Qubit Fluorometer (Qubit 4.0 Fluorometer, Thermo Fisher Scientific, USA) and the DNA was diluted to a final concentration of 3 ng L^-1^. Small aliquots were used to avoid freeze and thaw cycles. DNA extraction blanks were not concentrated, but used purely for subsequent PCR analyses.

For the amplification of fungal DNA, we used the tagged forward primer ITS67 and the tagged reverse primer 5.8S, developed for use on sedimentary DNA (Seeber et al., 2021). The amplified region has a size of approximately 183 bp. The use of tagged primers is essential to enable the assignment of the DNA sequences to the original samples after NGS sequencing. For each sample batch, six replicates were conducted independently from each other.

For the reconstruction of the palaeovegetation, we used the chloroplast trnL P6 loop marker region. The tagged primers being used are trnL g as the forward primer and trnL h as the reverse primer (Taberlet et al., 2007). For each sample batch, three replicates were conducted independently from each other.

A single PCR reaction contained in total 25 μL consisting of 3 μL DNA at a concentration of 3 ng μL^-1^, 0.2 μM of each primer, 10x HiFi buffer, 2 mM MgSO4, 0.1 mM dNTPs (consisting of dGTP, dATP, dCTP, and dTTP), 0.8 mg mL^−1^ BSA, and 1.25 U Platinum Taq High Fidelity DNA Polymerase (Invitrogen, United States). Each PCR batch also contained 3 μL of the corresponding DNA extraction blank and a PCR negative control with 3 μL of DEPC-water (deionised and diethylpyrocarbonate treated) instead of the DNA sample. All steps including DNA extraction and PCR set-up were conducted in the palaeogenetic laboratories at AWI Potsdam.

The PCR reaction itself was conducted in the Post-PCR laboratories at AWI Potsdam, which are located in a separate building to avoid contamination of ancient DNA samples with amplified DNA. The reaction for fungal ITS marker amplification was conducted in a thermocycler (Biometra, Germany) following the protocol for voucher samples for the samples from Lake CH12 (Seeber et al., 2021) while the other samples were amplified using the following protocol: starting with an initial denaturation at 94 °C for 2 min, followed by 40 cycles of 30 sec denaturation at 94 °C, 30 sec annealing at 54 °C, and 30 sec elongation at 72 °C, and a final elongation step of 10 min at 72 °C. The thermocycler protocol for plant trnL P6 loop amplification followed the protocol of Epp et al. (2018).

The PCR products were checked by gel electrophoresis (2% agarose gels). Only those products showing the expected gene bands were used for purification and subsequent sequencing. Purification was done with the MinElute PCR Purification Kit (Qiagen, Germany) according to the manufacturer’s protocol with the final elution in 50 μL. DNA concentration was measured with a Qubit Fluorometer (Qubit 4.0 Fluorometer, Thermo Fisher Scientific, USA). For sequencing, 40 ng of each purified PCR product were pooled. If no concentration was measurable, the total volume of the purified PCR product was added to the pool. For extraction blanks and PCR non-template controls, 5 μL of each PCR product was added. The final pool was purified again with a MinElute PCR Purification Kit (Qiagen, Germany) and adjusted to a final concentration of 33 ng μL^−1^ in a total volume of 30 μL. In total, three fungal sequencing pools were sent to Fasteris SA sequencing service (Switzerland). The service included the library preparation using a specified protocol (Metafast library) and a quality control and sequencing on an Illumina MiSeq platform (2 × 250 bp, V3 chemistry with an expected output of about 20 million paired-end reads).

We also sequenced two pools for the plant metabarcoding. Pooled purified PCR products of plant trnL P6 loop marker amplifications were sent to Fasteris SA together with other metabarcoding projects from our lab and sequenced on an Illumina NextSeq500 device (2 × 150 bp, 120 million paired-end reads). In addition, plant trnL P6 loop data from the lake CH12 were used from Epp et al. (2018).

### 3.3 Bioinformatic analysis

The subsequent analysis of the sequencing results was done using the open source OBITools pipeline (Boyer et al., 2016). A detailed description of all filtering steps can be found in Supplement 2. As a first step, *illuminapairedend* was conducted to pair the ends of the sequences, followed by *obigrep* to filter out only the joined sequences. Afterwards, *ngsfilter* was used to demultiplex the file into the unique samples and *obiuniq* was used to dereplicate the sequence reads. Then, all sequences shorter than 10 bp and with fewer than 10 reads were deleted applying *obigrep*. For the fungal dataset, the open source *sumaclust* algorithm (Mercier et al., 2013) was applied to cluster sequences with an identity threshold of 0.97 to generate operational taxonomic units (OTUs). After the filtering steps, *ecotag* was applied to perform the taxonomic classification of the OTUs against the embl142 (based on the EMBL nucleotide sequence database, release 142 (EMBL142; Kanz et al., 2005) and the UNITE database release for the fungal metabarcoding (Nilsson et al., 2019). The UNITE database is a curated fungus database where the detection of false positive reads might be lower than in the broader EMBL release. Using only the UNITE database for the assignment would potentially preclude identification of certain taxa. Therefore, the final assignment is based on the assignment from the database with the higher identity for each OTU. When both databases produced the same identity, the UNITE database was used for the final taxonomic classification.

For the taxonomic classification of the vegetation dataset, we used the ArctBorBryo database which is based on the quality-checked and curated Arctic and Boreal vascular plant and bryophyte reference libraries (Sønstebø et al., 2010; Willerslev et al., 2014; Soininen et al., 2015).

All databases were built after the following procedure to be applicable for the *ecotag* algorithm. The sequences of the databases and the NCBI taxonomy files were downloaded and both formatted in the ecoPCR format. Then, ecoPCR was run to simulate an *in silico* amplification of database sequences with the ITS67 and the 5.8S_fungi primers (allowing 5 mismatches in each primer sequence). The putatively amplified sequences were used as the reference databases and taxonomy information was added.

Resulting OTUs with identity levels equal to or higher than 98% were used for further analyses of the fungus dataset to keep only well annotated sequences. This is necessary to be certain about the analysis of fungus-plant relationships. For the analysis of the vegetation data, the identity cut-off was at 100%. PCR negative controls and extraction blanks were checked for contamination. All contaminants (non-fungal reads and OTUs occurring in non-template controls and extraction blanks) and aquatic fungi in the samples as well as OTUs with total read counts lower than 10 have been excluded from further analysis. Afterwards, we resampled both datasets to normalise the count data. The vegetation data have been resampled to a base count of 12,489 which resulted in an exclusion of the samples from 9.9 cal ka BP of Lake Kyutyunda and 7 cal ka BP from CH12 as they had too low counts. The fungus data were resampled to a base count of 5,284 resulting in a sample from 5 cal ka BP from Lake Kyutyunda and a sample from 18.8 cal ka BP from Lake Lama being excluded due to low read numbers.

### 3.4 Data analyses

Further filtering of the fungus dataset followed the suggested steps of Schiro et al. (2019). We assigned the identified fungus taxa to functional types according to their role in the ecosystem (Schulze and Mooney, 2012). The mycorrhizal fungi include arbuscular mycorrhizae, ectomycorrhiza, and ericoid mycorrhizae. The other groups are the saprotrophs, parasites, lichens, yeasts, and other symbionts besides mycorrhizae. A large number remained as “unknown” if their role in the ecosystem is not yet well understood. The identified plant taxa were assigned to either woody or herbaceous taxa.

All statistical analyses have been carried out using R, version 4.0.3 (R Core Team, 2020) using percentage data. The taxa were plotted colour-coded after their assigned functional type. Plotting has been done using the tidyverse package and ggplot2 in R (Wickham, 2016). To analyse differences in species diversity amongst the samples and locations, we calculated the alpha diversity using the function specnumber() of each sample from the resampled (rarefied) fungus and plant dataset.

To investigate the relationship between fungi and vegetation, we first assessed whether there is a correlation between fungal OTU richness and plant ASV richness. Second, we related the fungal richness to the most significant vegetation PCA axis scores. Finally, we applied the significant vegetation PCA axes as constraining variables in an RDA performed on fungal compositional data. For that, the scores of the PC axes were merged as a data frame. For each axis, only the taxa which make up most of the separation of the axes were plotted with their names in the final RDA to not overload the RDA with too many taxa. The significance of the vegetation PC axes was identified using PCAsignificance(). We used only 10 samples from Lake CH12 for the RDA to balance the weight of all lakes in the ordination. All ordination analyses were performed on double square-rooted data.

## 4. Results

### 4.1 Fungi: sedaDNA sequencing results and overall patterns of alpha diversity and taxonomic composition

In total, we obtained 52,213,129 counts in the fungal dataset. After assembling paired-end reads with *illuminapairedend*, demultiplexing into samples with *ngsfilter*, and cleaning the sequences with *obigrep* and *obiuniq*, we have 25,751 unique sequences with 32,027,606 counts. Clustering at a similarity threshold of 97% with *sumaclust* resulted in 5,411 OTUs. Excluding all OTUs with a lower similarity than 98% against the reference databases led to 716 remaining OTUs for the embl142 database, whereas the UNITE database returned only 268 different OTUs. After resampling to a base count of 5,284 and the subsequent filtering steps, 118 OTUs remained, covering 95.25% of the entire dataset, which was investigated. The other OTUs are regarded as “rare” and were not further assessed.

The highest OTU numbers before subsequently filtering taxa were detected in lake CH12 (209 OTUs). This is followed by Bolshoe Toko (146 OTUs) which is surrounded by forest and lake Levinson Lessing (137 OTUs) and lake Lama (135 OTUs). The lowest OTU number was detected for the northern lake Kyutyunda (78 OTUs). The OTU richness of single samples ranges from 3 OTUs (lake Kyutyunda, 30 cal ka BP) to 82 OTUs (CH12, 5.5 cal ka BP) with a mean of 23.53 OTUs. Samples from the Holocene show a higher richness in comparison to samples from MIS2 and MIS3.

The 10 most dominant taxa, which sum up to 71% in the entire fungal dataset, are Pseudeurotiaceae (20%; 30 samples), *Mortierella* (13%; 63 samples), Sordariomyceta (11%; 26 samples), *Exophiala* (5.8%; 6 samples), *Oidiodendron* (5.6%; 10 samples), *Protoventuria* (5.5%; 14 samples), *Candida vartiovaarae* (3.1%; 7 samples), *Pseudeurotium* (2.7%; 9 samples), *Gryganskiella fimbricystis* (2.6%; 32 samples), and *Trichosporiella cerebriformis* (2.4%; 11 samples).

The most dominant fungal functional type in the dataset is constituted by the saprotrophs (40%; 38 OTUs), while yeasts are present at 10% (23 OTUs). Parasites (9.05%; 13 OTUs) and mycorrhizae (4.5%; 14 OTUs) are relatively rare. The lowest abundances are the other symbionts (1.07%; 5 OTUs), lichens (0.2%; 4 OTUs), and mould (0.2%; 1 OTU). The fungi with unknown functions comprise 24.2% (21 OTUs) of the dataset.

### 4.2 Vegetation: sedaDNA sequencing results and overall patterns of alpha diversity and taxonomic composition

In total, we obtained 48,939,032 reads for the vegetation data. Assembling of the paired-end reads, demultiplexing into samples, and cleaning resulted in 152,194 reads. A total of 243 amplicon sequence variants (ASVs) were obtained with a 100% similarity to references in a curated reference library of arctic and boreal plants (Sonstebo et al., 2010; Willerslev et al., 2014; Soininen et al., 2015).

The comparison of the assigned sequences of the vegetation data between the lakes shows that lake Lama has the highest number of recovered and assigned sequences (163), followed by lake Bolshoe Toko (152) and lake Levinson Lessing (146). Lake CH12 (138) and lake Kyutyunda (133) have the lowest numbers of sequences assigned. The ASV richness of a single sample varies between 9 (7 cal ka BP, CH12) and 112 (35 cal ka BP, Bolshoe Toko).

The most common plant taxa are Salicaceae (37.4%; 69 samples), *Dryas* (20.4%; 69 samples), *Larix* (5.94%; 44 samples), *Alnus alnobetula* (5.88%; 67 samples), *Papaver* (3.86%; 59 samples), *Menyanthes trifoliata* (3.83%; 45 samples), *Bistorta vivipara* (2.72%, 63 samples), Asteraceae (2.43%; 66 samples), *Betula* (1.6%; 67 samples), and *Anemone patens* (1.4%; 18 samples). These ten taxa constitute 85.5% of the whole dataset.

### 4.3 Site-specific plant-fungus co-variation

#### 4.3.1 Fungus and plant co-variation in Arctic Siberia from MIS3 to the Holocene

In the Levinson Lessing record (northern Taymyr Peninsula, tundra, 40–0 cal ka BP), the Pseudeurotiaceae as well as *Mortierella* and *Gryganskiella fimbricystis* (both saprotrophs) are highly abundant during MIS3 (Figure 2). Around 38 cal ka BP, the Didymellaceae (parasitic fungus family) also occur. At the end of MIS3, *Thamnolia vemicularis* (lichen) occurs. The most abundant plant taxa at this time are Salicaceae, *Dryas*, and *Papaver*. The most abundant fungus taxa in MIS2 are also Pseudeurotiaceae (unknown function) and *Mortierella* (saprotroph), but *Trichosporiella cerebriformis* (unknown function) also occurs often. For the plants, the most dominant taxa are Salicaceae and *Papaver*, followed by *Dryas* at the end of MIS2 (Figure 3). During the Holocene, *Mortierella* remains the most frequent fungal taxon but more mycorrhizal OTUs (*Inosperma calamistratum, I. geraniodorum, Mallocybe fuscomarginata, Oidiodendron*) and parasites (Didymellaceae, *Kalmusia variispora*) become abundant as well. In the Holocene, there is a drastic decline in *Papaver* while *Alnus alnobetula* becomes highly abundant at the beginning of the Holocene. *Dryas* as well as Salicaceae remain mostly abundant.

**Fig. 2:**
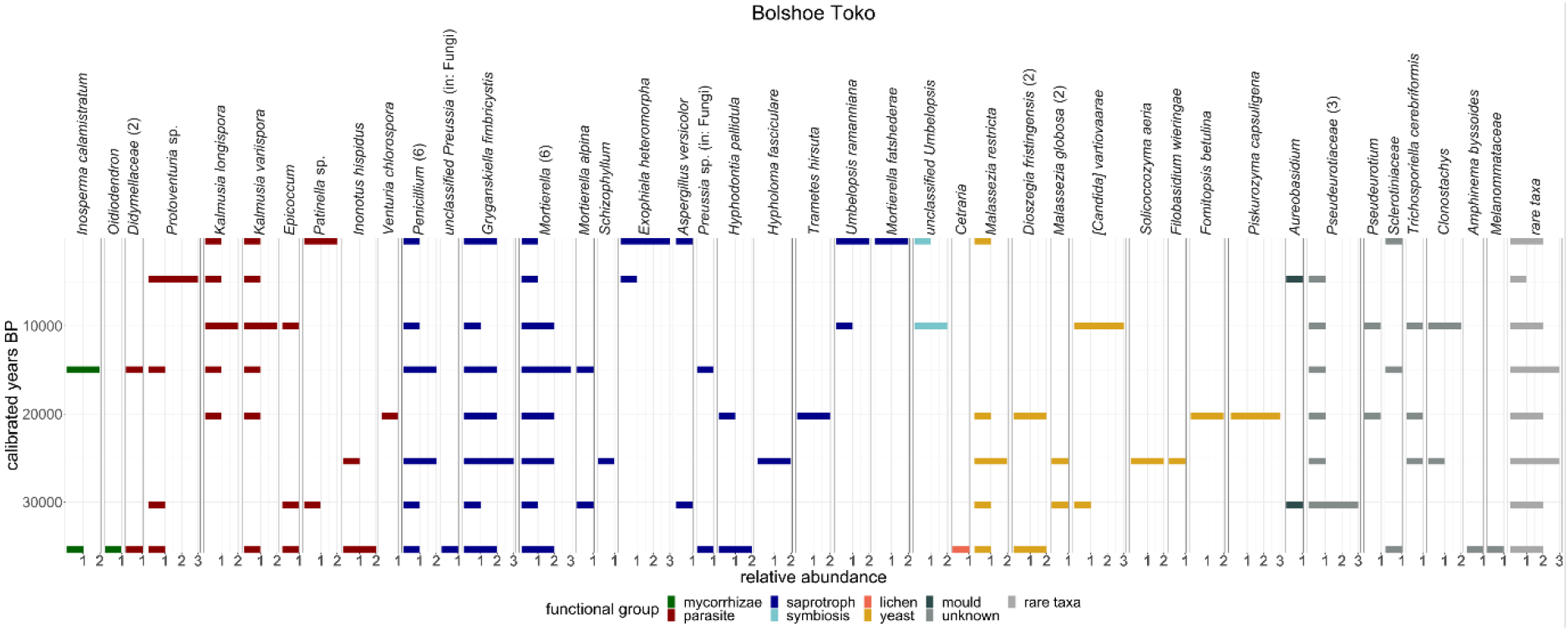
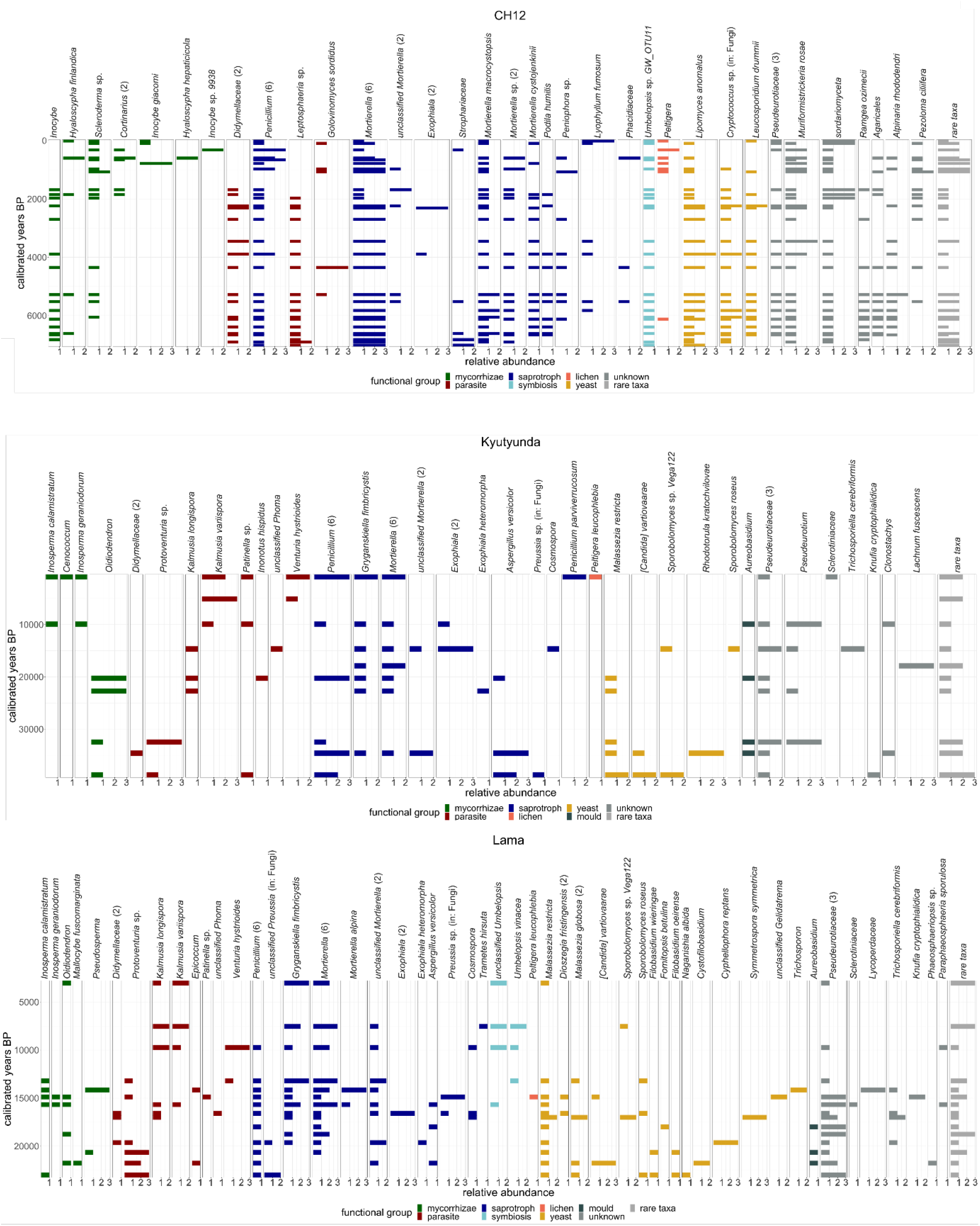
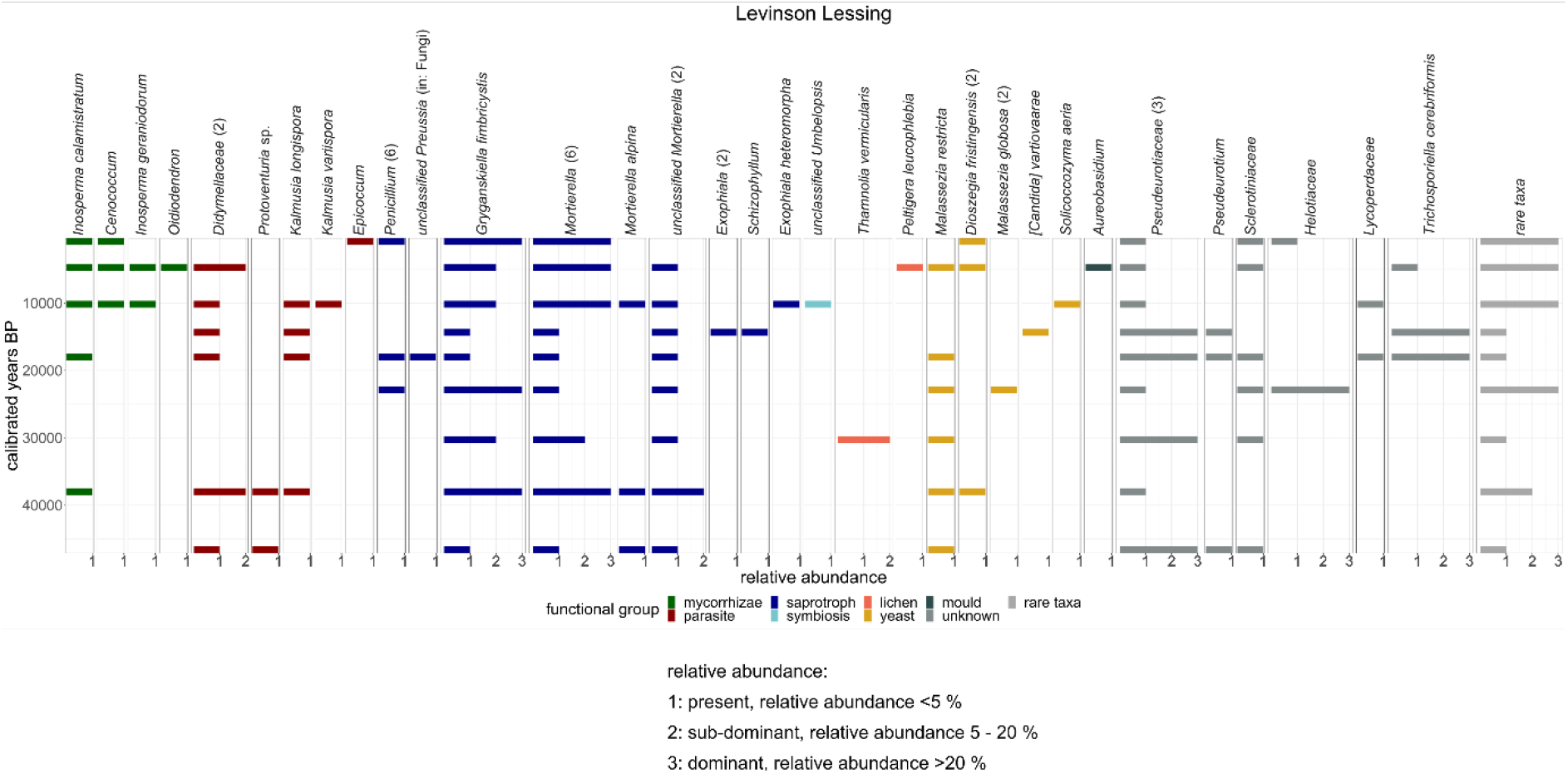
Site-specific fungus abundance displayed in relative percentages. Fungi of the same functional type are colour-coded. The numbers in brackets give the OTUs detected for the specific taxon.

**Fig. 3:**
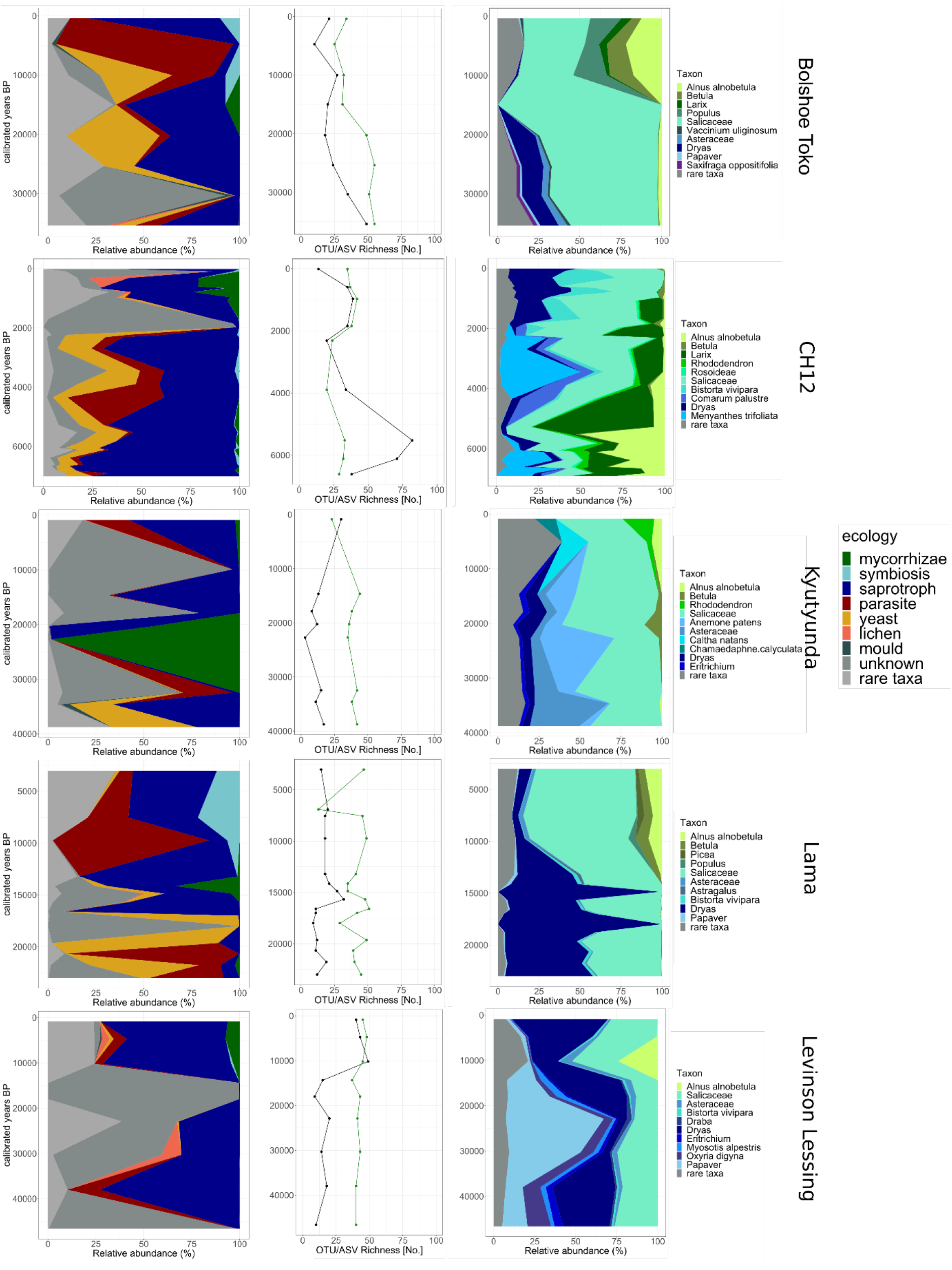
Fungal functional types in relation to fungal OTU richness and dominant plant taxa. Left column: distribution of fungal functional types for each lake. Middle column: fungal OTU richness of each lake (total OTU numbers), with the black line representing the fungal taxa while the green line marks the vegetation ASVs as comparison. Right column: ten most dominant plant taxa of each lake.

The lake CH12 record (southern Taymyr Peninsula, tundra, 7–0 cal ka BP) only spans the mid to late Holocene. Around 7 cal ka BP, *Inocybe* (mycorrhizae) as well as *Golovinomyces sordidus* and Didymellaceae (parasites) are highly abundant. *Mortierella* is present throughout the whole record but shows strong declines when mycorrhizae and parasites are abundant around 5 cal ka BP (Figure 2). Until 5.5 cal ka BP, this lake shows high abundances of *Alnus alnobetula* and Salicaceae. Woody taxa such as *Alnus alnobetula, Larix, Betula*, and *Rhododendron* have their highest abundances around 5 cal ka BP (Figure 3). After 5 cal ka BP, an increase in yeast taxa (e.g. *Lipomyces anomalus, Cryptococcus*) is detected. This coincides with a decline in the aforementioned woody taxa. In this record, the lichen genus *Peltigera* is abundant in more recent times when the variety of mycorrhizal taxa also increases and *Inocybe, Hyaloscypha finlandica, Scleroderma, Cortinarius, Inocybe giacomi*, and *Hyaloscypha hepaticola* occur. Saprotrophic taxa such as *Mortierella* species, *Lyophyllum fumosum, Penicillium*, and *Exophiala* are present throughout the whole record.

The record of lake Lama (northern Siberia, tundra-taiga transition zones, 24–0 cal ka BP) spans MIS2 and the Holocene. The most abundant fungal taxa during MIS2 are Pseudeurotiaceae, *Protoventuria* (parasite), *Mortierella* (saprotroph), and *Cyphellophora reptans* (yeast) (Figure 2). *Dryas* as well as Salicaceae dominate the vegetation. Around the beginning of the Bølling/Allerød (15 cal ka BP), *Pseudosperma* and *Inosperma* species (mycorrhizae) become abundant. A little later, *Venturia hystrioides* and *Kalmusia* species (all parasites) start to occur. Salicaceae is still the most dominant plant taxon, but *Alnus alnobetula, Picea, Betula*, and *Populus* frequently occur after 15 cal ka BP as well. Additionally, a drastic decline in *Dryas* took place after 15 cal ka BP (Figure 3).

Lake Kyutyunda covers the late MIS3 to the Holocene (northern Siberia, tundra-taiga transition zones, 38.8–0 cal ka BP). The most dominant fungal taxa during the late MIS3 are *Oidiodendron* (mycorrhizae), *Pseudeurotium* (unknown function), and *Penicillium* (saprotroph) (Figure 2). During this time, Salicaceae and Asteraceae are the most abundant plant taxa but *Alnus alnobetula* also occurs occasionally (Figure 3). *Oidiodendron* is mainly present at the end of MIS3. Shortly after, a large increase in *Betula* is detectable. In MIS2, the fungal taxa *Oidiodendron* and *Penicillium* are still highly prevalent and the taxon *Lachnum fuscescens* (unknown function) becomes common (Figure 2). Salicaceae remain the most dominant plant taxon and *Betula* starts to occur more frequently. High abundance of *Dryas* as well as the first instances of *Alnus alnobetula* are detectable in the late MIS2 (Figure 3). During the Holocene, *Pseudeurotium* (unknown function) and *Kalmusia variispora* (parasite) became the most abundant fungal taxa. Salicaceae maintained its broad distribution while other woody taxa such as *Alnus alnobetula* and *Rhododendron* increased in their abundances.

The samples from Bolshoe Toko also span the late MIS3 to the Holocene (central Yakutia, taiga, 35–0 cal ka BP). During the late MIS3, the Pseudeurotiaceae are the most abundant fungal family in this lake but parasitic species (e.g. *Kalmusia* species, *Inonotus hispidus*) and saprotrophs (e.g. *Mortierella, Grykanskiella fimbricystis*) also occur (Figure 2). At this time, Salicaceae has the highest abundance amongst the plants with *Dryas* occurring frequently. *Alnus alnobetula* and *Betula* are also present but at low abundance (Figure 3). In MIS2, *Gryganskiella fimbricystis* and *Mortierella* are highly abundant fungi and a few yeast taxa (e.g. *Dioszegia fristringensis, Piskurozyma capsuligena*) start to occur. In the late MIS2, *Inosperma calamistratum* (mycorrhizae) also occurs. The vegetation is still dominated by Salicaceae until the end of MIS2 with scarce abundances of *Alnus alnobetula* and *Betula*. In the Holocene, *Protoventuria* (parasite) is the most abundant fungal taxon but also *Kalmusia* species (parasite), *Exophiala heteromorpha* (saprotroph), and *Candida vartiovaarea* (yeast) are commonly found. A large increase in more diverse woody taxa is detected with more occurrences of Salicaceae as well as *Alnus alnobetula, Betula, Larix*, and *Populus*.

#### 4.3.2 Quantitative relationships between fungi and plant richness and composition

Taking all records together, we found only a weak borderline-significant correlation between fungi OTU and plant ASV richness (*r* 0.2394, *p*-value 0.098). Fungal richness is positively correlated to the samples scores of the first plant PCA axis (PC1: *r* 0.3863, *p*-value 0.006) and negatively correlated to the sample scores of the second plant PCA axis (PC2: *r* -0.41, *p*-value 0.003). The first axis reflects the differences between samples characterised by woody taxa including *Larix* and *Alnus alnobetula* and typical tundra taxa. On the second axis, we detected herbaceous plant taxa such as *Anemone patens* and *Thymus* positively correlating alongside other taxa preferring wetter habitats. Taxa such as *Oxyria digina* and *Dryas*, which are associated with rather dry sites, show a negative correlation.

Sample scores of plant PCA axes 1–5 explain 20% of fungi composition (*p* 0.001) as revealed by RDA (Figure 4). The PCA of the vegetation composition can be found in Supplementary 3. Woody taxa such as *Alnus alnobetula, Larix*, and *Rhododendron* appear in the upper right quadrant of the RDA plot together with the fungal taxa *Mortierella*_003 (saprotroph), *Cryptococcus* (yeast), and *Muriformistrickeria rosae* (unknown function) (Figure 3) and samples from CH12 aged 5.5 and 1.8 cal ka BP. The RDA also shows that parasitic fungi, such as Didymellaceae, and yeast, such as *Lipomyces anomalus* and *Cryptococcus*, tend to occur in the presence of woody taxa. Lichens occur predominantly in samples of Holocene age. *Papaver* and *Dryas* together with the fungal taxa Pseudeurotiaceae, *Grykanskiella fimbricystis* (saprotroph), and *Mortierella* (saprotroph) species occur in the lower left quadrant together with all samples from Levinson Lessing. *Populus* and *Ranunculus* in the upper left quadrant appear together with *Lipomyces anomalus* (yeast), *Cryptococcus* (yeast), and *Mortierella* (saprotroph). The samples here mostly originate from Bolshoe Toko although there are also some from lake Lama (around the Bølling/Allerød period) and lake Kyutyunda (Holocene). The samples from the Holocene all occur in the upper half of the RDA where the woody plant taxa are found and a broader fungal species richness is detected. In general, Lake CH12 shows a unique fungal composition in comparison to the other lakes. The samples can be found in the right quadrants of the RDA while the samples of the other lakes are located in the left quadrants or centred (Figure 4).

**Fig. 4:**
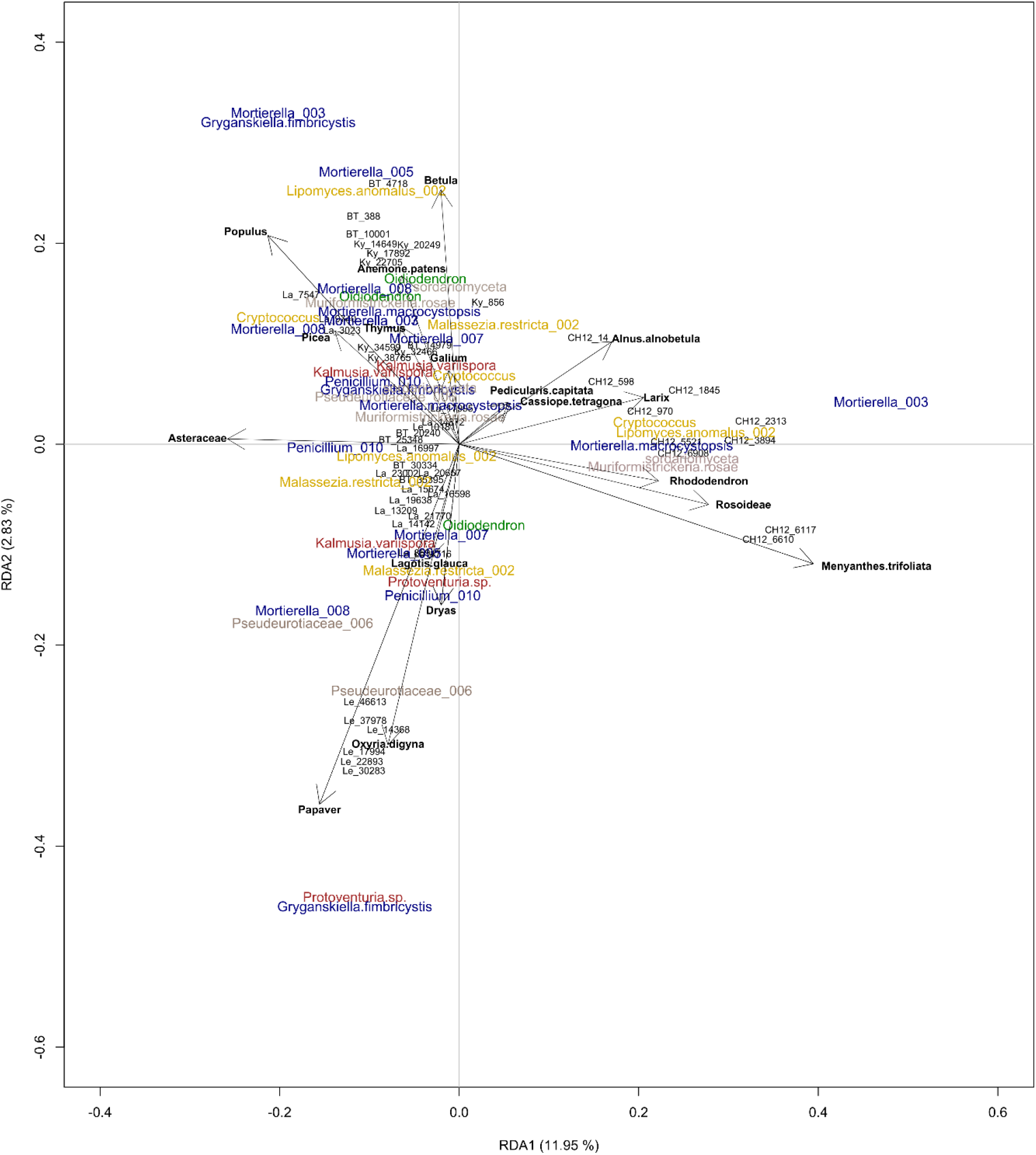
Fungal and plant co-variation displayed in a redundancy analysis (RDA). The most relevant principal component axes of the vegetation were determined, the scores extracted and then integrated into the RDA. The fungal taxa are displayed colour-coded according to their functional group (see Figures 2 and 3). The plant taxa are marked with black arrows. The numbers after the taxa names indicate the specific OTU. The sample names are shortened with the lake name (BT= Bolshoe Toko, Ky = Kyutyunda, La = Lama, Le = Levinson Lessing) and the calibrated year BP. The vegetation explains 20% of the fungus distribution.

## 5. Discussion

### 5.1 Fungi richness along vegetation gradients in Siberia

Using sedimentary DNA metabarcoding of 70 samples from five lake sediment cores from Siberian tundra and forested sites, we tracked a high fungal richness (706 OTUs). So far, Siberia has not been targeted by high-throughput sequencing analyses focusing on fungi diversity over a spatial gradient despite hundreds of these studies being available worldwide (Baldrian et al., 2021). In our study, we used OTU clusters instead of observed species, which could have led to an under- or overestimation of richness (Frøslev et al., 2017). Underestimation of richness may originate from missing reference material of arctic and alpine taxa in databases including species that have not yet been described (Goodwin et al., 2016; Quince et al., 2009). A species assessment from the western Ural Mountains, covering a similar environmental setting, yielded a richness of a similar order of magnitude, that is of 376 different fungal species (Palamarchuk and Kirillov, 2019). This supports the conclusion of Seeber et al. (2021) that the marker is suitable to assess fungal diversity even on a very long time-scale. For comparison, higher OTU richness (1125 OTUs in 55 samples) was obtained by Talas et al. (2021) in their study of a Holocene lake sediment core from eastern Latvia. The higher overall diversity is explained by the inclusion and detection of aquatic fungi (about 23% of their taxa, terrestrial about 40%), which are mostly lacking in our data. In addition, Talas et al. (2021) did not remove very short reads and included reads with fewer counts (4 instead of 10 in our study).

Lake Bolshoe Toko (146 OTUs) and lake Lama (135 OTUs) from forested areas show overall higher OTU richness in comparison to lake Kyutyunda (78 OTUs) from the northern tundra (Figure 3). A relationship between fungal species richness and vegetation composition is shown by many studies (e.g. Tedersoo et al., 2013; Geml et al., 2017), however studies from the Siberian treeline ecotone are hitherto lacking. Our study agrees with the finding that ectomycorrhizal fungal richness is highest with forest cover (Geml et al., 2017). The spatial fungal richness gradient is confirmed by the temporal relationship – we observed a co-occurrence of high fungal richness and woody vegetation (as inferred from sedaDNA using the trnL P6 loop marker). For example, the lake Levinson Lessing record shows a large increase in the fungal OTU richness and woody taxa dominance during the warm Holocene compared with the (late) Glacial period. In contrast, experimental warming did not result in higher fungal diversity (Geml et al., 2015; Mundra et al., 2016). The lake Lielais Svētiņu sediment study from the Holocene by Talas et al. (2021) showed high richness as well as community turnover with increases in plankton parasitic species and mycorrhiza after 4 cal ka BP, suggesting that those fungi that are more specific with their hosts or substrates – such as ectomycorrhizae – are more susceptible to ecosystem changes than taxa with wide preferences. In our data, lake CH12 shows higher OTU richness than the other lakes, even when a similar number of samples is considered, which supports the hypothesis that fungal communities from the warmer Holocene might be more species rich. Potentially, warming-induced vegetation responses rather than direct warming shape the diversity in fungal communities. This suggests that future treeline shifts northwards will bring along a broadening diversity of fungi species.

A study from sub-arctic Canada observed an overall higher fungal diversity on moss tissue compared with conifer litter (Matsuoka et al., 2021) indicating that in addition to the overall vegetation gradient, further biotic (He et al., 2017) and abiotic (Genevieve et al., 2019) factors including soil condition and wetness are important (Geml et al., 2016). Metabarcoding studies on arctic tundra communities reveal that even for each specific tundra type, the communities of the associated fungi are unique (Wallenstein et al., 2007; Geml et al., 2021). This might explain the overall highest fungal OTU richness originating from lake CH12 (a dry forest tundra site), while lake Kyutyunda (a wet southern tundra site) shows a rather low richness. Furthermore, our analysis shows a negative correlation between fungal richness and the second vegetation PC axis, covering a wetness gradient from species related to drier areas (high PC scores) to species rather related to wetter areas.

A study on multiple sites of the Tibetan plateau showed that fungal richness is positively correlated with plant richness (Yang et al., 2017). Interestingly, we find only a weak positive correlation between fungal richness and plant richness, which might originate from a complex relationship between plant richness and vegetation composition. Also, we assessed plant ASVs and fungal OTUs which makes a direct comparison difficult. Incomplete databases for arctic fungi might also lead to underestimation of taxa. It is known that at the broad scale, plant richness decreases with latitude (Kerkhoff et al., 2014). However, a study of modern vegetation from Kamchatka, Russian Far East, reports highest plant species richness in alpine tundra and snowbed communities (Doležal et al., 2013). A recent sedaDNA study from the treeline zone in Chukotka (Russian Far East) and from Bolshoe Toko (which is also part of the present study) showed that terrestrial plant richness was particularly high in the late Pleistocene in steppe-tundra areas and lower in the forested Holocene (Huang et al., 2020; Courtin et al., 2021). This indicates that a high correlation between fungus and plant alpha diversity cannot be expected.

### 5.2 Changes in fungal functional groups related to vegetation transition

#### 5.2.1 Mycorrhiza

Most mycorrhizal taxa detected and most reads obtained all over the analysed cores are from the families Cortinariaceae and Inocybaceae and some from Myxotrichaceae and Hyaloscyphaceae (Figure 2). This is in accordance with previous fungal metabarcoding studies (Nilsson et al., 2005; McGuire et al., 2013; Botnen et al., 2014). We observed *Inocybe* (including the subgenus *Inosperma* (Matheny et al., 2020)) and *Cortinarius*, both of which are known from high latitudes (Timling et al., 2012). They represent ectomycorrhiza associates of arctic tundra and shrubs including *Salix* and *Dryas integrifolia* (Ryberg et al., 2009; Botnen et al., 2014), which are common taxa in our plant metabarcoding dataset. It also supports previous studies from boreal forests (McGuire et al., 2013), including a study from the Russian Far East which detected *Cortinarius*, in addition to *Lactarius* and *Russula*, as important ectomycorrhiza of *Larix gmelinii* in this region (Miyamoto et al., 2021). In addition to Pinaceae, *Cortinarius* associates with shrubs of Salicaceae and Rosaceae as well as herbaceous Cyperaceae (Garnica et al., 2005), which are common families in our plant metabarcoding dataset. Nevertheless, mycorrhizal fungi in the treeline area are highly dependent on the specific locality itself and its soil properties such as pH, nutrient availability, and C:N ratio (Toljander et al., 2006).

It has been reported earlier that, as a consequence of the Last Glacial, vegetation species richness decreased as well as arbuscular mycorrhizal taxa while ectomycorrhizal and non-mycorrhizal fungi increased (Zobel et al., 2018). This resulted in changes in the mutualist trait structure after the Last Glacial Maximum (LGM), leading to an increase of ectomycorrhiza associated with woody taxa and a decrease in arbuscular mycorrhiza forbs. Therefore, mycorrhizal associations are important factors when predicting the species responses to changing environmental conditions (Zobel et al., 2018). We also observe a generally increasing fungal richness in the Holocene samples (Figure 3). Long-term changes in the diversity of fungal communities are highly dependent on soil age, and richness increases with growing soil age (Cutler et al., 2014). This strengthens our observations and underlines the suitability of sedaDNA fungal metabarcoding studies for appropriate ecosystem reconstructions.

Interestingly, we observed highest values of Pinaceae only after the presence of mycorrhizal taxa (Cortinariaceae and Inocybaceae; e.g. *Cortinarius* and *Inosperma calamistratum*) (Figures 2 and 3), although this might be by chance because of our low sample numbers. If no mycorrhizal fungi are present in a habitat, growth of Pinaceae individuals is slowed down or establishment is inhibited as nutrient uptake is not possible for non-mycorrhizal roots (Marschner and Dell 1994). Studies from Japan (Ishida et al., 2007) and temperate areas in the Himalaya (Pande et al., 2004) revealed Cortinariaceae as one of the main ectomycorrhizal associates of Pinaceae, which strengthens the preciseness of our dataset and its possibility to correctly recover fungal-plant covariation over a long time scale. Our analysis also highlights the longevity of the dependency of the Pinaceae family on these particular fungi.

#### 5.2.2 Saprotrophs

We found *Mortierella, Penicillium*, and *Exophiala* species as the main biomass-decaying taxa (Figure 2). These are common soil fungi also in high-latitude ecosystems (Treseder et al., 2007; Allison et al., 2009). *Mortierella* and *Penicillium*, two of the most common taxa in our dataset, are reported as some of the main soil fungi in arctic tundra soils (Kurek et al., 2007; Zhang et al., 2016) due to their cold tolerance. *Mortierella* was also found as an associate of *Vaccinium uliginosum, Betula nana, Salix glauca, Empetrum nigrum*, and *Cassiope tetragona* (Voříšková et al., 2019), which are typical taxa in our metabarcoding study. Rhizosphere samples from *Larix sibirica* and *Betula pendula* from Krasnoyarsk, Siberia revealed *Penicillium* as one of the main constituents (Boyandin et al., 2012). On the other hand, we did not find typical saprotrophs known from central Siberian forests unaffected by permafrost such as *Fomitopsis pinicola, Hymenochaete cruenta, Rhodofomes cajanderi*, and *Trichaptum abietinum* (Park et al., 2020). Nevertheless, *Larix* forests growing on permafrost show a broad host spectrum towards saprotrophic species (Leski and Rudawska, 2012) as a response to changing environment, for example after wildfires (Miyamoto et al., 2021), which favours their survival and explains why different studies recover other associated taxa.

Saprotrophs are generally highly abundant throughout all records. The significant decrease in saprotrophic fungi around 10 cal ka BP for all lakes (Figures 2 and 3) demonstrates that the drastic climate change during the Pleistocene/Holocene transition (Biskaborn et al., 2016; 2021) also affected soil communities. This finding from natural past warming agrees with results from experimental warming studies, indicating that relative saprotroph abundance declines with warming, while the abundance of mycorrhizal fungi and lichen increases (Deslippe et al., 2012; Geml et al., 2015; Mundra et al., 2016).

#### 5.2.3 Parasites

The most abundant parasitic species from our dataset are *Protoventuria, Kalmusia variispora, Kalmusia longispora*, and Didymellaceae which co-occur with Salicaceae, *Larix*, and *Alnus alnobetula* (Figures 2 and 3). *Protoventuria* species, for example, penetrate leaves and show up as distinct spots on the plant leaf (Carris and Poole, 1993). In shrubby tundra in Greenland with *Salix* occurrences, *Venturia* species are amongst the highest abundant fungi (Voříšková et al., 2019), indicating a strong covariation between this fungus and *Salix*. To date, plant-parasite interplay in relation to climate change is not fully understood (Burdon and Zhan, 2020) but it is assumed that parasitic fungal species are more specific in their hosts than, for example, mycorrhizal taxa are (Põlme et al., 2018).

Generally, we detected parasitic OTUs mostly in sediment samples from the warm Holocene (Figure 2). This confirms early findings that experimental warming leads to an increase in parasitic and virulent fungi (Geml et al., 2015) along with woody taxa expansion. Interestingly, we can observe a drastic decline in Salicaceae after the high *Protoventuria* abundance around 20 cal ka BP (Figures 2 and 3), which supports previously recovered high fungal parasite abundances in permafrost during the last ice age (Lydolph et al., 2005). Fungi from the family Venturiaceae have been assigned to Salicaceae as pathogenic species in northern latitudes (Hosseini-Nasabnia et al., 2016), while *Kalmusia* was detected in *Alnus* forests in Lithuania (Iznova and Rukšėnienė, 2012). Didymellaceae is co-occurring with a broad range of host plants such as *Larix decidua* (Chen et al., 2017). In our data, the RDA reveals that *Kalmusia* species preferentially occur in forested areas alongside saprotrophic and mycorrhizal species (Figure 4). This supports the value of our data and the feasibility of co-occurrence analysis in sedaDNA studies.

#### 5.2.4 Lichens

Our analyses are among the few palaeo-ecological studies that have detected lichens (Figure 2). They are commonly lacking in fossil records (Taylor and Osborn, 1996) despite being an important component of boreal forest and tundra biomass (Asplund and Wardle, 2017; Shevtsova et al., 2020). However, we could only detect very few lichen reads, belonging to 48 OTUs (less than 1% of the whole dataset). Generally, environmental metabarcoding has been assessed as a valuable tool to investigate modern lichen communities (Fernández-Mendoza et al., 2017) and yielded higher OTU richness than the analysis of voucher specimens from the same region (Wright et al., 2019). Lichen DNA has been reported to drastically degrade after a few hundred years (Kistenich et al., 2019). This means that lichens might be harder to recover from broad fungal palaeo-metabarcoding studies than other functional groups. A specific set of markers established for the amplification of lichen DNA targeting not only the fungus part of the lichen but also the phyco- and cyanobionts could be an asset to improve the taxonomic resolution as well as to trace diversity.

In total, the recovered lichen OTUs belong to 16 families. The families with the highest OTU numbers are Peltigeraceae and Parmeliaceae and the most abundant genera are *Thamnolia, Peltigera*, and *Cetraria* with all of them being common elements in northern Siberian communities (Zhurbenko and Yakovchenko, 2014) and permafrost (Lydolph et al., 2005). *Thamnolia* species often occur in arctic tundra communities (Sheard, 1977) and are of low specificity concerning their photobiont as they associate with various *Trebouxia* species (Nelsen and Gargas, 2009). *Peltigera* preferentially grows in temperate regions on soils and among mosses over rocks, but can also be found on tree trunks (Nash 2002) and in boreal forests (Asplund and Wardle, 2015), explaining their abundance in the more forested Holocene in our records from CH12 (Figure 2).

Interestingly, almost all lichens are recorded from sediments of warm periods with well-developed vegetation, namely the late MIS3 and the Holocene. This is to some extent unexpected, as lichens are a prominent feature of arctic landscapes and short-term experimental warming in the Canadian arctic led to a decline in lichen abundance (Fraser et al. 2014). For Siberia, lichens have been recovered along a broad latitude gradient with high diversity and biomass (Safronova and Yurkovsksya, 2019). Lichens have been discovered to possess a cooling effect when growing on permafrost, giving them great importance when considering thawing effects on permafrost (Porada et al., 2016). The implication of a potentially reduced or missing lichen cover during the glacial might be of relevance for past permafrost soil carbon dynamics. Lichens are reported to tolerate high percentages of CO_2_ (Badger et al., 1993), but studies about the impact of low CO_2_ supply are not available. Possibly, lichens suffer more than other fungi from reduced atmospheric CO_2_ content as they also have to supply their algal or cyanobacterial symbiont.

The distribution of lichens also has an impact on the occurrence of animals such as *Moschus moschiferus*, which preferentially settle in lichen-rich habitats for their food supply (Slaght et al., 2019). Increased lichen coverage during the Holocene may have supported the compositional turnover in the megaherbivore fauna. Reindeer mostly feed on lichen but changing environmental conditions will probably impact their distribution as well as their diet to include less lichen (Drucker et al., 2011) or seasonal variations in their diet (Bocherens et al., 2015), which would give them a higher survival advantage. Changing fungus communities will thus not only impact the boreal forest taxa, but also its fauna and general soil properties.

#### 5.2.5 Yeast

The most abundant yeast taxa from our dataset are *[Candida] vartiovaarae, Malassezia restricta, Cyphellophora reptans, Cryptococcus*, and *Lipomyces anomalus* (Figure 2). These taxa are widely distributed in soils in Siberia (Polyakova and Chernov, 2001). A study from Germany showed *[Candida] vartiovaarae* to be broadly present in forest as well as in grassland soils (Yurkov et al., 2012), while *Cryptococcus* is associated with peatland (Thormann, 2006) and boreal swamps (Kachalkin and Yurkov, 2012). A correlation between *Malassezia* species and soil nematodes in central European forests has been found, suggesting that the nematodes act as vectors for the small fungi (Renker et al., 2003). To investigate these zoophilic relationships for the palaeoenvironment, further metabarcoding data on small soil organisms could be an asset.

Whenever yeasts are highly abundant in the records, mycorrhizal taxa decrease (Figures 2 and 3) (and vice-versa making them antagonists in their response to warming and further environmental changes). In our dataset, yeasts were present preferentially in colder time periods like the LGM. Most yeast species can show an adaptive response when temperatures drop to maintain their survival (Kandror et al., 2004). Aside, experimental warming studies underline our finding that yeast taxa decline with rising temperatures (Treseder et al., 2016). This indicates that some yeast species will lose their habitats with ongoing warming, resulting in a major feedback on the ecosystem, potentially leading to ecosystem turnover.

From our data, it is not possible to determine the role of yeast in the soil. Soil yeast can serve both as biotoxins (Santos et al., 2004; Compant et al. 2005) or growth promoters for plants (Nassar et al., 2005; El-Tarabily and Sivasithamparam, 2006). It is likely that in Siberian soils, yeasts either function as plant parasites (Hernández-Fernández et al., 2021) or as biodegraders, as after a period of high yeast abundance, we detect a decrease in woody taxa. A better understanding of modern mutualistic and parasitic interactions in Siberian tundra and taiga soils will help to solve this research gap.

### 5.3 Implications of our results for ecosystem functioning and future research avenues

As fungi are a key component for ecosystem functioning, a major impact on future ecosystem-climate feedbacks can be expected arising from fungi compositional change in concert with changes of the whole soil microbiome in permafrost (McCalley et al., 2014). So far, the complex interplay between climate, vegetation, fungi, and microorganisms in the boreal forest ecosystem is not yet understood. To our knowledge, we conducted the first study on fungus-plant interactions and co-occurrences in the palaeo context, assessing community shifts in boreal forests as well as tundra ecosystems. However, our results can only be seen as a first proxy on future fungal community changes in response to warming as the magnitude of warming differs strongly between the studied samples and present warming and any relationship may incorporate lagged responses on decadal to millennial time-scales (Biskaborn et al., 2021).

To our knowledge, this is the first long-term dataset to show this antagonistic relationship among the fungal functional types. Our study shows that warming-related vegetation change is relatable to fungus diversity and fungal functional changes. By analogy to the past, woody taxa advancing into arctic regions in the future will result in a higher fungus diversity and a relative increase in mycorrhizae, parasites, and potentially lichens at the cost of saprotroph and yeast abundance.

Our study design does not allow a definite conclusion to be drawn as to whether or not future treeline advances might rely on the presence of specific fungal communities. As ectomycorrhizal communities in the sub-arctic tundra are generally species-rich and do not show a high host preference (Ryberg et al., 2009; 2011), major changes may not be expected. However, the investigated soils in the sub-arctic already have a long history of soil development, which does not apply to bare northern tundra sites and upper mountain areas. These are potential habitats for forest establishment from a temperature point of view but might not be favourable for a diverse soil fungus composition due to a lack of nutrients and lower temperatures.

By analogy to the results of our study, lichens do not generally suffer from warming but can be affected by the vegetation. The observed decline in lichens as a consequence of denser canopy cover (Cornelissen et al., 2001) may only apply to the more southerly forests. As our study only returned a few lichen OTUs, it is not possible to draw a robust conclusion here. Extensive CO_2_-concentration supply during experimental darkening leads to a generally quick accumulative CO_2_ uptake in, for example, the genus *Peltigera* and a subsequent relatively slow release (Badger et al., 1993), making lichens potentially valuable for the storage of future warming-induced CO_2_ release from soil. Further research into lichens could be promising for the development of mechanisms to support ecosystem adaptations towards changing environments.

Besides the limitation in the temporal resolution, our study suffers from limited taxonomic resolution and a complex abundance pattern. Sedimentary ancient DNA metabarcoding is a highly complex method which faces several limitations. First, it is highly susceptible to any damage and degradation which can lead to biases in the PCR products as potential taxa might be dismissed due to their short lengths (Coissac et al., 2012; Taberlet et al., 2012). Second, sometimes the reference genomes are missing and therefore identification at the relevant taxonomic resolution is not possible (Sønstebø et al., 2010). Third, different taxa possess varying amounts of copy numbers per cell which can also lead to an overrepresentation of taxa with high copy numbers while rare taxa might be missed (Behnke et al., 2011).

## 6. Conclusions

This is the first study showing spatial and temporal changes in palaeo fungus-plant covariation. Knowing which fungi influenced the growth of specific plant communities in the past will help to predict future community turnover due to varying climate. To understand palaeo community turnover in more detail, it is necessary to consider a plant’s associated heterotrophic organisms in present times. This will help to place the knowledge gained in this study into better context. Additionally, our data are a great asset to the existing knowledge about boreal forests as they help to shed light on adaptation mechanisms of plants towards warming and their subsequent northward migration. Nevertheless, there are still many ecological interactions unknown which need to be addressed in future research, such as which organisms contribute to the rhizospheres of specific plants and if or how these associations change with varying climate. Despite this, our findings will already help the assessment of future tipping points in boreal forest stability.

## Declarations

### Availability of data and material

The data are available under xxxx and will be publicly available after the acceptance of the manuscript.

Pangaea: Metadata of the cores and links to existing Pangaea entries

Dryad: Fungal DNA Dataset (Raw data and scripts)

Dryad: plant DNA Dataset (Raw data and scripts)

### Author contribution

The study was designed by UH, KSL, BvH; KSL supervised and BvH conducted the experimental lab work; BvH analysed the data under supervision of UH and KSL; LS sampled the cores and supervised the DNA extractions; PS and LE performed the bioinformatic evaluation of the marker; MM retrieved the sediment core of Lake Lama; BD led the projects on Kyutyunda and Bolshoe Toko; BB retrieved and dated the sediment cores including age depth modelling of Bolshoe Toko, Kyutyunda, CH12 and Levinson Lessing; BvH dated Lama and performed the age modelling; BvH supervised by UH wrote a first version of the manuscript; all authors commented on the first and revised version of the manuscript.

