## Supplementary for "Long-term fungus–plant co-variation from multi-site sedimentary ancient DNA metabarcoding in Siberia"

**Supplementary 1:** Available radiocarbon ages (years before 1950 CE) from Biskaborn et al. (2016) next to slightly corrected composite depths, dating error and method applied in the Poznan radiocarbon laboratory. RES insoluble humin fraction; SOL alkali-soluble humic acids fraction; TOC total organic carbon. To perform age-depth modelling we used RES values from bulk sediment samples.

| 14C Lab ID | Sample ID | 14C age (yrs) | 14C error (yrs) | Depth below sediment surface (cm) | Sample type | Method |
|---|---|---|---|---|---|---|
| Poz-49481 | PG2023-2_50,5-51 | 2405 | 35 | 76,75 | bulk | RES |
| Poz-49472 | PG2023-2_71,5-72 | 3585 | 30 | 93,25 | bulk | RES |
| Poz-49471 | PG2023-2_188-188,5 | 5900 | 40 | 210,25 | bulk | RES |
| Poz-49470 | PG2023-3_79-79,5 | 8000 | 50 | 304,75 | bulk | RES |
| Poz-49474 | PG2023-3_187-187,5 | 9420 | 50 | 412,75 | bulk | RES |
| Poz-49483 | PG2023-3_244-244,5 | 10620 | 60 | 469,75 | bulk | RES |
| Poz-49482 | PG2023-4_78,5-79 | 13360 | 100 | 492,75 | bulk | RES |
| Poz-50559- | PG2023-5_25-26 | 16870 | 260 | 638 | bulk | SOL |
| Poz-50560 | PG2023-5_25-26 | 27820 | 300 | 638 | bulk | RES |
| Poz-50557 | PG2023-4_231-232 | 29180 | 350 | 644 | bulk | RES |
| Poz-50558- | PG2023-5_25-26 | 29990 | 380 | 638 | bulk | TOC |
| Poz-50555- | PG2023-4_231-232 | 33500 | 500 | 644 | bulk | TOC |
| Poz-49484 | PG2023-5_162,5-163 | 33770 | 350 | 775,25 | bulk | RES |
| Poz-50556- | PG2023-4_231-232 | 33900 | 700 | 644 | bulk | SOL |

**Supplementary 2:** Bacon age-depth model based on radiocarbon age determinations (bulk sediment samples, RES values, Supplementary 1) from sediment core PG2023 retrieved in 2010 from Lake Kytyunda. This is a refined version of the age-depth correlation for this sediment core published by Biskaborn et al. (2016).

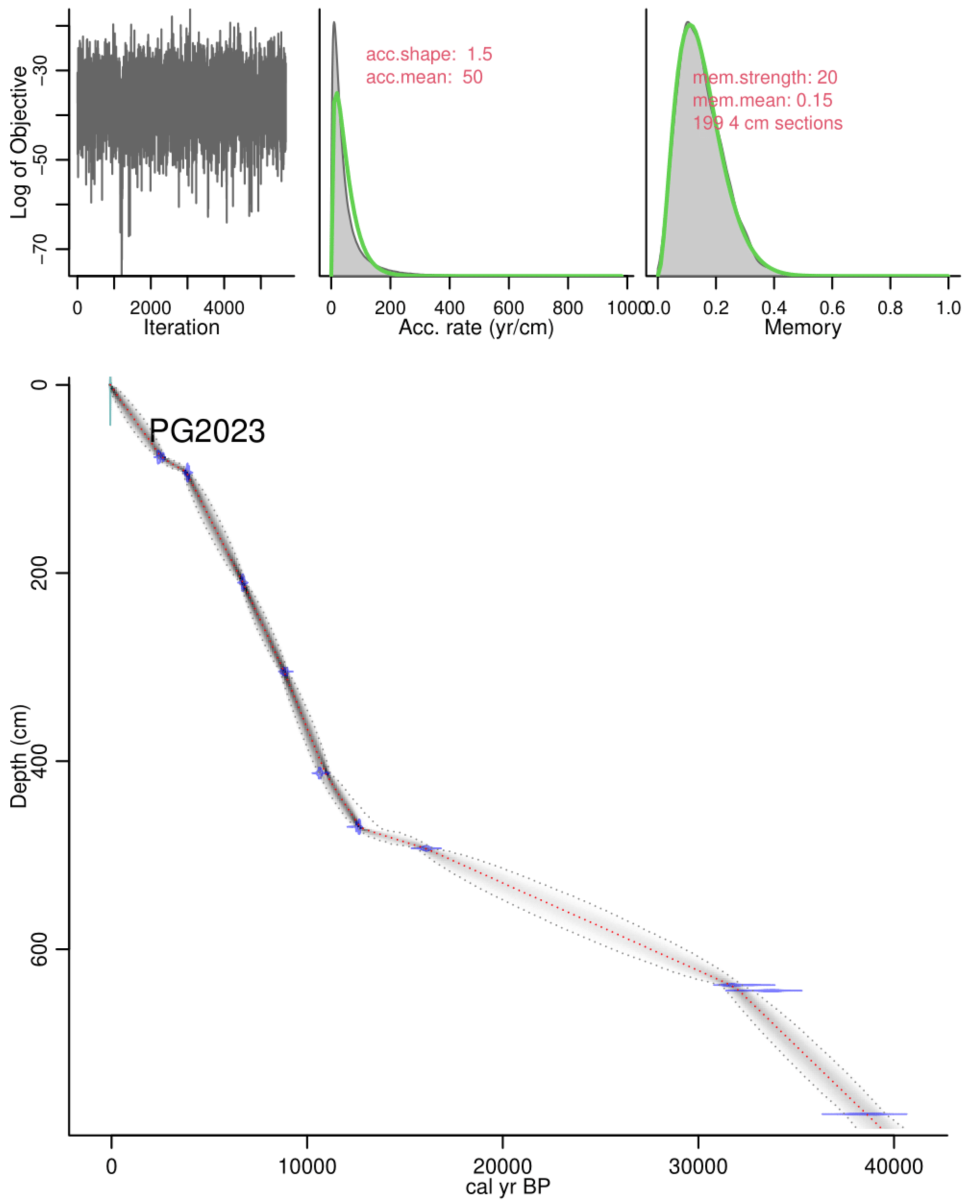

**Supplementary 3:** Radiocarbon ages from the Lama PG1341 core which were used to calculate the age-depth model alongside dated woody remains (*: data from Andreev et al. (2014)). The reservoir effect for this core is 4460 years and was subtracted from the determined 14C ages before the age-depth modelling.

| 14C Lab ID | Sample ID | 14C age (yrs) | 14C error (yrs) | Depth below sediment surface (cm) | Sample type | Method |
|---|---|---|---|---|---|---|
| 6794 | PG1341-4AR_15-16 | 4597 | 25 | 15-16 | Bulk | C14 |
| 6795 | PG1341-4AR_42-43 | 4845 | 25 | 42-43 | Bulk | C14 |
| 6796 | PG1341-4AR_80-81 | 5698 | 25 | 80-81 | Bulk | C14 |
| 6797 | PG1341-4AR_105-106 | 6530 | 26 | 105-106 | Bulk | C14 |
| 6036 | PG1341-4_140 | 7333 | 62 | 140-141 | Bulk | C14 |
| UTC8876 | Woody remains | 5255* | 48 | 211 | Woody remains | C14 |
| UTC8877 | Woody remains | 6200* | 60 | 253 | Woody remains | C14 |
| 6037 | PG1341-5_260 | 11258 | 83 | 260-261 | Bulk | C14 |
| 6038 | PG1341-5_380 | 13365 | 100 | 380-381 | Bulk | C14 |
| 6039 | PG1341-5_503 | 17536 | 154 | 503-504 | Bulk | C14 |
| 6040 | PG1341-6_620 | 17400 | 104 | 620-621 | Bulk | C14 |
| 6041 | PG1341-6_740 | 16954 | 101 | 740-741 | Bulk | C14 |
| 6042 | PG1341-7_860 | 17480 | 105 | 860-861 | Bulk | C14 |
| 6043 | PG1341-7_980 | 18594 | 120 | 980-981 | Bulk | C14 |
| 6044 | PG1341-8_1046 | 18887 | 397 | 1046-1047 | Bulk | C14 |
| 6045 | PG1341-8_1220 | 18011 | 354 | 1220-1221 | Bulk | C14 |
| 6046 | PG1341-9_1340 | 18976 | 395 | 1340-1341 | Bulk | C14 |
| 6047 | PG1341-9_1460 | 18967 | 119 | 1460-1461 | Bulk | C14 |
| 6048 | PG1341-10_1580 | 20900 | 501 | 1580-1581 | Bulk | C14 |
| 6049 | PG1341-10_1700 | 21296 | 531 | 1700-1701 | Bulk | C14 |
| 6050 | PG1341-11_1820 | 23707 | 683 | 1820-1821 | Bulk | C14 |

**Supplementary 4**: Age-depth model for Lake Lama. The age-depth model was established using the package bacon() in R. The reservoir effect is 4460 years. We used two dated woody remains alongside 19 radiocarbon-dated bulk sediment samples to set up the age model. A change in the sedimentation rate after a depth of 500 cm is visible, leading to a hiatus in the age model.

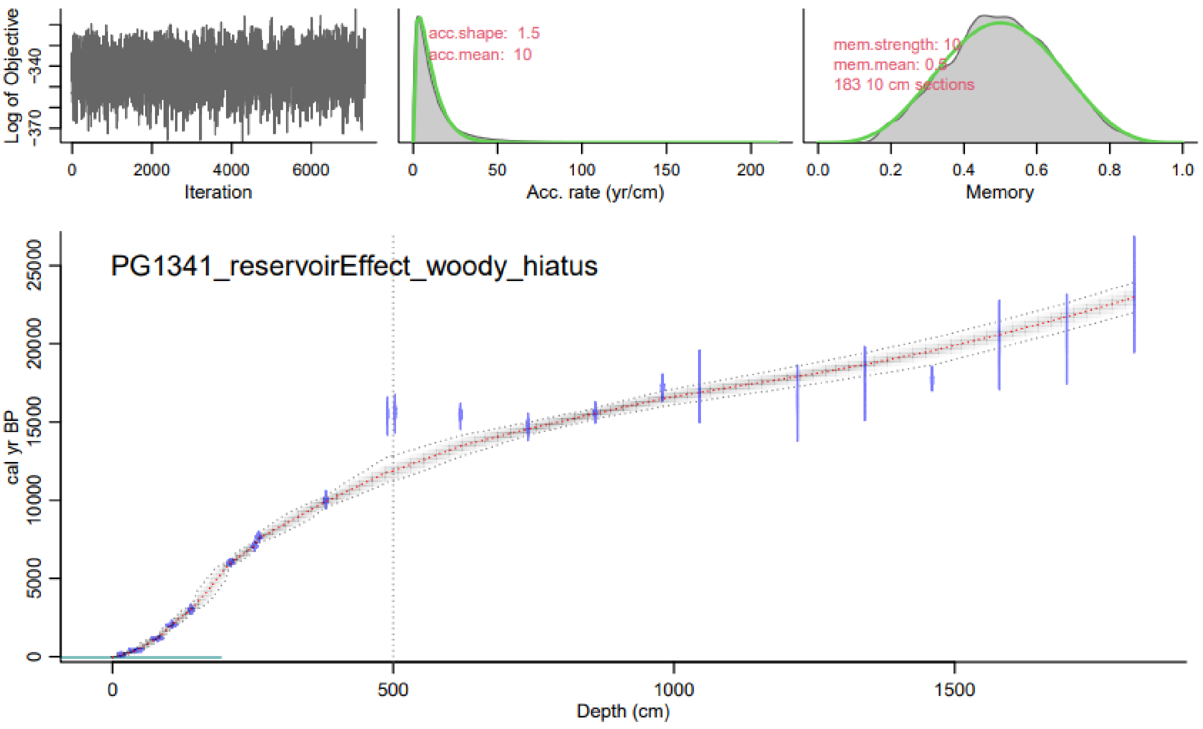

**Supplementary 5**: Obitools Pipeline and Cleaning Steps. The single steps and their commands are listed in the table as well as an explanation of each step. The last column includes the resulting size of the fungal data as an example.

| **Step** | **Command** | **Result (in brackets: the size of the fungal dataset after the subsequent step)** |
|---|---|---|
| illuminapairedend | illuminapairedend inputR1.fastq -r inputR2.fastq > paired_end.fastq | Paired-ending of sequences (52,213,129) |
| obigrep | obigrep -p ‘mode!="joined"' paired_end.fastq > paired_end_joined.fastq | Greps out only joined sequences (52,213,129) |
| ngsfilter | ngsfilter -t tagfile.txt -u unident.fastq pairedend_joined.fastq > assigned.fastq | Demultiplexing into samples (40,699,830) |
| obiuniq | obiuniq -m sample assigned.fastq > assigned_unique.fastq | Dereplicate sequence reads (3,294,811) |
| obigrep | obigrep -l 10 -p 'count>=10' assigned_unique.fastq > assigned_unique_l10_c10.fastq | Delete sequences with length shorter than 10 and counts lower than 10 |
| obiclean | obiclean -s merged_sample -r 0.05 -H assigned_unique_l10_c10.fastq > assigned_unique_l10_c10_clean.fastq | Selects sequences according to head, singleton, and interval; if a sequence is less than 20 times more frequent it will be assigned to the sequence with the higher count under the requirement that the difference is -d number of differences between the sequences (default 1), if you use option -H head sequences will be selected only (26,665; 151,151) |
| sumaclust (only for ITS dataset) | ./sumaclust -t 0.97 assigned_unique_l10_c10_clean.fastq > assigned_merged_datasets_sumaclust97.fasta | Clustering of sequences into OTUs using a similarity threshold of 97% between cluster centres and member sequences (5411 cluster created) |
| obigrep (only after sumaclust) | obigrep -p 'cluster_center' assigned_merged_datasets_sumaclust97.fasta > assigned_merged_datasets_sumaclust97_centres.fasta | Extraction of the cluster centres |
| ecotag | ecotag -R database.fasta -d database assigned_merged_datasets_sumaclust97_centres.fasta > | Taxonomic assignment of the OTUs with the database (either UNITE, embl or ArctBryo) |

| | | |
|---|---|---|
| | assigned_merged_datasets_sumaclust97_centres_database.fasta | |
| obiannotate | obiannotate --delete-tag=explain assigned_merged_datasets_sumaclust97_centres_database.fasta > assigned_merged_datasets_sumaclust97_centres_database_anno.fasta | Adds sequence record annotations, deletes the attribute named "explain" |
| obitab | obitab -o assigned_merged_datasets_sumaclust97_centres_database_anno.fasta > assigned_merged_datasets_sumaclust97_centres_database_anno.txt | Change fasta to tabular format |
